# Temporally unconstrained decoding reveals consistent but time-varying stages of stimulus processing

**DOI:** 10.1101/260943

**Authors:** Diego Vidaurre, Nicholas E. Myers, Mark Stokes, Anna C. Nobre, Mark W. Woolrich

## Abstract

In this paper, we propose a method to track trial-specific neural dynamics of stimulus processing and decision making with high temporal precision. By applying this novel method to a perceptual template-matching task, we tracked representational brain states associated with the cascade of neural processing, from early sensory areas to higher-order areas that are involved in integration and decision-making. We address a major limitation of the traditional decoding approach: that it relies on consistent timing of these processes over trials. Using a temporally unconstrained decoding analysis approach, we found that the timing of the cognitive processes involved in perceptual judgements can vary considerably over trials. This revealed that the sequence of processing states was consistent for all subjects and trials, even when the timing of these states varied. Furthermore, we found that the specific timing of states on each trial was related to the quality of performance over trials.

## Introduction

Neural processing of a stimulus and its use in guiding behaviour are highly dynamic. A given stimulus typically elicits a cascade of activation across the brain, including its multiple parallel as well as re-entrant pathways, starting with early feature analysis and leading to increasing integration and decision making (see, for example, Meyers et al., 2008, Harvey et al., 2012). But, to which extent is it possible to capture the progressive stages of this information-processing cascade related to stimulus processing from non-invasive human brain imaging data?

An influential approach has been to use decoding models that capture how the current stimulus is represented in the brain activity (Norman et al., 2006; Haynes and Rees, 2006; Tong et al., 2012; Haxby et al., 2014; Grootswagers et al., 2017). Assuming we have a number of trials or repetitions of a certain process (e.g. the presentation of a stimulus), the standard approach for decoding is to separately train one classifier or regression model (depending on whether the stimulus is categorical or continuous) at each time point, by pooling together the data from all trials in the training set, and then testing for the accuracy of each of these models on a test data set (King and Dehaene, 2014). This can then be used to interrogate the temporal dynamics of the processes evoked by the stimulus (see e.g. Meyers et al., 2008; Carlson et al., 2011; Isik et al., 2013; Carlson et al., 2013; Stokes et al., 2013; King et al., 2014). However this approach is, by construction, based on the assumption that these brain processes are *synchronous* across trials, i.e. the different stages of information processing start and finish at the same time within each trial.

Here, we argue that assuming consistent timing over trials may be too restrictive, ignoring trial-to-trial variability in the dynamics. Furthermore, it can severely misrepresent the data by leading to a potentially false conclusion of persistent activity, artefactually created by the act of averaging across trials (Stokes and Spaak, 2016; Latimer et al., 2015; Lundqvist et al., 2016). Sometimes, this assumption can also induce the impression of having a high number and relatively rapid succession of distinct processing states, as a consequence of trial-to-trial variability in the onset and duration of a potentially much smaller succession of states. Such temporal variability of processing states could be ubiquitous. For example, different stages of information processing may start and finish at different time points in each trial, depending on different levels of arousal or selective attention at the time of stimulus onset, or as a result of learning and plasticity.

In this paper, we propose a new framework based on the Hidden Markov model (Rabiner, 1989) and Bayesian variational inference (Vidaurre et al., 2016), for identifying representational brain states with high temporal resolution and with no assumption about the states having to occur at fixed time points on each trial. We refer to it as Temporally Unconstrained Decoding Analysis (TUDA). A functional, representational state is defined here as a decoding model that characterises a relation between brain activity and the current stimulus (Haynes and Rees, 2006). Each state is distinct from other states in describing how and in which regions the brain represents the stimulus, such that a switch of state indicates that the decoding does not cross-generalise before and after the switch (i.e. the stimulus-specific pattern of activity has changed). TUDA thus estimates the decoding weights associated to each decoding model, and identifies when each decoding model is “active” for each trial. Crucially, this is done without restricting the model to be active at the same point in time on each trial.

By applying this approach to magnetoencephalography (MEG) recordings in a perceptual judgement task, we found that, when allowing for this temporal flexibility, a reduced number of decoding models (fewer than six) is sufficient to explain the between-trial temporal differences in the data. This compares with standard decoding, which, with one model per within-trial time point (typically more than 100), cannot access this information at all. Furthermore, we found that the temporal dynamics of the decoding models correlate with behavioural changes over trials, lending additional support to the physiological relevance of between-trial temporal variability of the underlying neural processing cascade. To be able to meaningfully relate this information to behaviour is not only useful, but also proves the existence of tangible and interpretable differences in stimulus processing between trials.

## Results

Decoding analysis (**Figure 1a**) is a popular tool for interrogating the temporal dynamics of stimuli representation in the brain. Here, we propose a new method, Temporally Unconstrained Decoding Analysis (TUDA), which finds decoding models, or states, that characterise how the current stimulus is represented in the brain at different points in time. Crucially, TUDA estimates the exact timing of the decoding models together with the decoding weights that define each decoding model. This approach requires many fewer parameters to predict the stimulus from the data than the standard decoding approach, which assumes consistent timing over trials, and fits one model per time point (**Figure 1b**). In exchange, TUDA adds a new degree of freedom: that is, *when* each decoding model is active in each trial, which is inferred in a data-driven manner.

**Figure 1.**
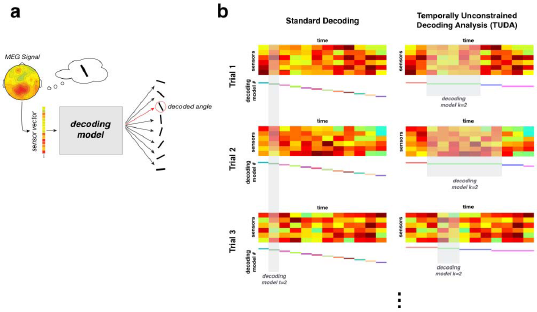
**(a)** General representation of decoding analysis. **(b)** A schematic representation of the standard decoding approach (left) vs. the Temporally Unconstrained Decoding Analysis (TUDA) (right).

Here, we apply TUDA to sensor-space MEG data collected when subjects were performing a perceptual judgement task, in which they were shown an oriented visual grating stimulus and asked to compare it to a memorised template orientation to detect matches (the difference between these being referred to as the *relative* angle). The model was estimated separately for each participant (of which there were ten) and each session (of which there were two per participant), obtaining a set of decoding models and model time courses indicating the probability of each model being active at each time point within each trial. We estimated the model for different numbers of decoding models (*K*=3,4,5,6). More details about the method and the experimental paradigm are presented in the Methods section.

### Standard decoding misrepresents the data if trials are not synchronous

In order to show how standard decoding can misrepresent the data when stimulus processing is not synchronous across trials, we tested TUDA and the standard decoding approach on synthetic data. In this scenario, we generated synthetic data using three decoding models (each, a linear function mapping data to stimulus) that cycle through during the trial; that is, all trials start with decoding model 1, dwell some time until switching to model 2, then move to model 3, come back to model 1, and so on. Given a 300Hz sampling frequency and 1s trials, the dwell time varies between 0.066s (20 time points) and 0.166s (50 time points) for each decoding model visit and each trial. In this way, the between-trial variability regarding which model is active increases as we progress through the trial. Specific details on the nature of the simulations can be found in the SI.

**Figure 2a** illustrates the performance of TUDA on these simulated data (see the SI also for a complete description of the results). The difficulties for the standard decoding approach in ignoring between-trial variability (and by using a larger number of decoding models than it exists) are illustrated in **Figure 2b,c**. We computed the Pearson correlation between sets of regression coefficients for each pair of models, and grouped the models according to their similarity. The results are shown in **Figure 2b**, where we can see a large fragmentation into several different models. If we now assess the performance of each decoding model on held-out trials we can obtain an across-time generalisation matrix (King and Dehaene, 2014). This is shown in **Figure 2c**, using cross-validated explained variance (CV-R2) as the summary statistic. It can be observed that the trials generalise sharply at the beginning of the trials, since decoding model 1 is always active at start. Then, the pattern becomes blurrier as a consequence of the increasing between-trial temporal variability. Standard interpretations of this result would artefactually suggest that brain activity gains in persistency (generalisation) at the end of the trials, when, in reality, the models’ dwell time is the same across the entire trial.

**Figure 2.**
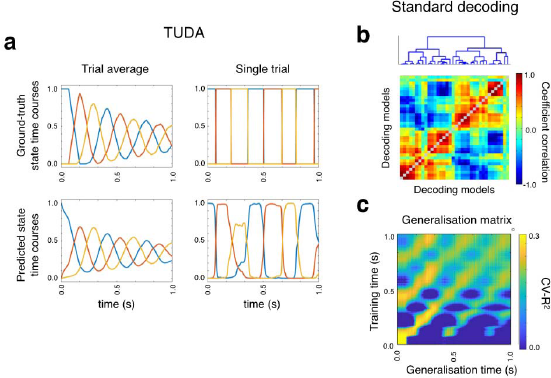
Failing to account for between-trial differences can result in a misleading view of stimulus processing. **(a)** True and predicted model time courses (on average and for a single trial) from the TUDA inference process on synthetic data with between-trial temporal differences. **(b)** Hierarchical clustering of the decoding models (using the correlation between their regression coefficients) suggests that the standard decoding approach (which estimates one model per time point) finds many distinct models (more than three, which is the actual number of models in the synthetic data). **(c)** Cross-validation showing the generalisation over time of the standard decoding approach (King and Dehaene, 2014) suggests a continuous and relatively fast fluctuation of models when looking at the diagonal of the matrix, which may lead to the incorrect conclusion that there are many different, non-exchangeable models underlying the data.

### The neural processes relevant to the task are not synchronous across trials

Using a model with *K*=5 decoding states, **Figure 3a** shows when each of the five inferred decoding models (represented using a different colour) is active as a function of time. This is shown for a subset of the trials ordered by reaction time (RT). Note that the decoding is, by necessity, carried out separately for each session, and so here we are presenting the results for a single participant and session. However, these results are representative of the results across the whole dataset (see SI). Underneath, **Figure 3b** shows the average occupancy (that is, the mean over all trials) of each decoding model as a function of time. We next show the extent of temporal variability of the decoding models for the proposed approach, i.e. when we do not constrain the trials to have the same decoding dynamics. For this purpose, we chose one of the decoding models that had a clear peak in the average occupancy time course, and took all trials where the model was active at the time of maximum model occupancy (represented by the marked peak in the bottom panel). In **Figure 3c**, histograms for the starting and finishing times of these model occurrences reveal large temporal variability.

**Figure 3.**
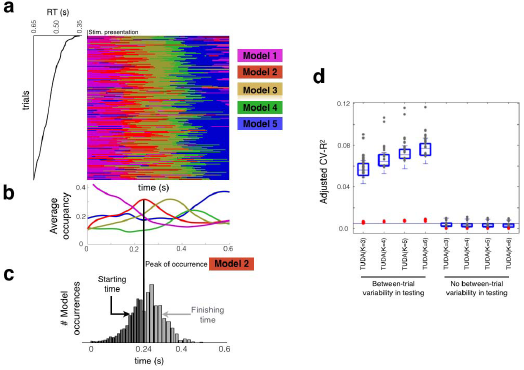
Decoding of the relative angle exhibits large temporal variability between trials. **(a)** Time-by-trial representation of which decoding model is active at each time point shown for a subset of the trials from a representative session and participant. Models are numbered according to the order in which they tend to arise in trials, and trials are ordered by reaction time (RT); only 200 trials are shown. **(b)** Percentage of trials (taken over all subjects and trials) assigned to each decoding model as a function of time. **(c)** For a given decoding model, and this specific session, temporal variability represented as a histogram of starting and finishing times for those trials when the chosen decoding model is active at time *t* (*t*=0.23s for presented angle and *t*=0.24s for relative angle). **(d)** TUDA’s accuracy, measured as adjusted cross-validated explained variance (adjusted CV-R2, see Methods), when accounting for between-trial variability vs. when between-trial differences are ignored, for different number of decoding models (*K*=3, 4, 5 and 6). Each grey dot represents one session, the red dots represent the baseline accuracy obtained from surrogate data (see Methods), and the horizontal line represents (group-level) cross-validated accuracy for the standard decoding approach.

We next investigated the extent to which accounting for between-trial temporal differences impacts the prediction accuracy of the model. Normally this would be estimated using cross-validation. However, TUDA uses both the data *and* the stimulus in combination to estimate the decoding model time-courses (see Methods). This means that the stimulus information for a held-out trial gets used to estimate the decoding model time course for that trial (as well as for predicting the stimulus). We correct for this bias through the use of surrogate data, to calculate an *Adjusted CV-R^2^* measure (see Methods), allowing us to compare between different TUDA approaches and standard decoding.

**Figure 3d** shows *Adjusted* CV-R2 for TUDA when we model between-trial temporal differences and when we do not, for different numbers of decoding models *(K)*. As a reference, the horizontal line represents CV-R2 for standard decoding. The accuracy when modelling between-trial temporal differences is orders of magnitude larger than when ignoring such differences, highlighting the importance of these differences, and is also superior to the standard decoding approach despite using fewer models.

Finally, if between-trial temporal variability is such an important factor, we argue that, when ignoring this variability, the TUDA predictions should be significantly worse for those time points with greater diversity in decoding model allocation. For *K*=5, **Figure SI-1a** shows, for the same illustrative session used before, CV-R^2^ as a function of time for the traditional decoding approach and TUDA. The peaks of accuracy closely correspond to the peaks of activation of the decoding models as shown underneath (where there are less between-trials temporal variability). Using all sessions, **Figure SI-1b** shows, for each time-point (represented as a dot), trial variability in the assignment of a decoding model in training (measured as the variance of the model time courses across trials) versus estimation accuracy in testing (measured using CV-R^2^); the Pearson correlation coefficient is 0.47 (p-value <0.0001, permutation testing).

### Sequences of states and their relation to behaviour

If between-trial temporal variability has a neural origin, then we might also expect it to relate to behaviour. We next analysed reaction time (RT) data, which was collected for all trials where the subjects pressed the button (i.e. when participants judge that the presented angle matched the template angle in their working memory). We discarded all trials with no button press. Importantly, we first regressed the absolute relative angle out of the model time courses and the RTs. This is necessary because RT could have a direct correlation with the relative angle: smaller relative angles might make participants more confident, leading to faster responses. Without this precaution, a relation between the model time courses and RT could be trivially driven by the actual relative angle, instead of the intrinsic between-trial variability that we are interested in.

We then estimated, for each decoding model, the correlation between the corresponding (deconfounded) model time courses and RT across trials (i.e. the Pearson correlation between the probability of the state being active and RT). For an example session, **Figure 4a** shows the resulting time-resolved correlations for each of the decoding models, reflecting a strong relation between the decoding models’ trial-specific timings and behaviour. On these grounds, we next examined how short RT trials compare to long RT trials, by viewing the temporal profile of which decoding model best predicts RT at each point in time. For an example session, **Figure 4b** shows these temporal profiles for prototypical *short RT* and *long RT* trials. The temporal profiles were calculated as follows: at each time point, the decoding model chosen to be active is the model with the highest across-trials correlation (for the short RT trial), or the highest anticorrelation (for the long RT trial), between the estimated probability of being active and RT. As can be seen, the same ordering of decoding models underlies *short RT* and *long RT* trials; however, the timing is very different, with the decoding models getting active around 0.25s earlier in the *short RT* as compared to the *long RT* trials. As illustrated in **Figure 4c**, this characteristic sequence is also (separately) found in the transition probability matrix between the decoding models, which reflects the estimated probability of transitioning between every pair of decoding models, and is inferred without knowledge of RT as part of the model inference (see Methods). This strong sequential order of the decoding models is largely present in all participants and sessions (**Figure SI-2**).

**Figure 4.**
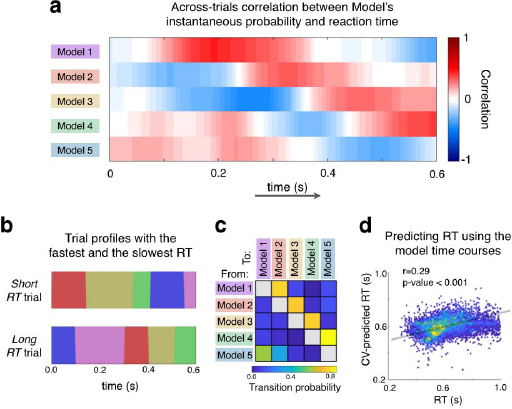
The precise timing of the decoding models within trials has an intimate relationship with reaction time (RT). **(a)** For the same representative session used in **Figure 2**, the correlations between RT and each decoding models’ activation probabilities as a function of time, are very high. **(b)** Prototypical sequences of decoding models for a *short RT* trial (top) and *long RT* trial (bottom) for the representative session. **(c)** The transition probability matrix for the representative session, containing the probability of transitioning between each pair of models, has a strong sequential structure (see Figure SI-4 for other sessions). **(d)** Cross-validated prediction of RT as using the model time courses confirms the strong relationship between the models’ temporal-variability and RT (each data point corresponds to one trial, and colour indicates density of points). The plot depicts all sessions, although the prediction was performed session by session.

We further evaluated the strength of the relationship between the model time dynamics and behaviour by predicting, in a cross-validation setting, the trial RT using the model time courses (after regressing out the absolute relative angle from both variables; see Methods for details). The prediction was done at the group level, i.e. using all participants together (cross-validation folds were constructed such that the entire set of trials of each subject were assigned to a single fold; Winkler et al., 2015). **Figure 4d** shows real versus predicted RT, where each dot represents a trial. The prediction accuracy is highly significant (p-value <0.001, permutation testing), confirming that the temporal decoding variability effectively relates to behaviour.

### Decoding models are spatially localised

We next examine the spatial characteristics of the decoding states. Interpretation of decoding weights is not straightforward (Weichwald et al., 2015), so we computed the *encoding* model that corresponds to each decoding model. For each sensor, decoding model and session, the encoding model is defined as the regression weights that predict the data for this sensor as a function of the relative angle, using only the time points when the decoding model is active (that is, making use of the model time courses; see Methods). **Figure 5** shows a summary of the spatial characteristics of the encoding models, which can be compared for reference with the maps shown in Myers et al. (2015). The topographic map represents the sum of explained variances across encoding models. Although there are differences across models and subjects, the maps indicate that the relative angle is encoded in motor and frontal sensors.

**Figure 5.**
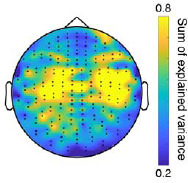
Topographical map in sensor space reflecting the (averaged) spatial activation associated to the estimated decoding models. This is expressed as the sum of explained variance (CV-R^2^) of the corresponding encoding models (see Methods).

## Methods

### Task and participants

This study used previously published data used for a different purpose (described in Myers et al., 2015). Ethical approval for methods and procedures was obtained from the Central University Research Ethics Committee of the University of Oxford. In brief, we recorded MEG data while participants performed a template-matching visual task (EEG data were also simultaneously acquired but were not used in the current study). Ten right-handed volunteers (age range: 21-27yrs, 6 females) took part in the study, completing two sessions each, containing short blocks. In each block, participants were presented with one orientation template to keep in mind. They then viewed a stream of oriented gratings, and responded when the presented angle matched the template angle.

The task consisted of eight brief (approximately six-minute) blocks, in which 480 stimuli were presented (resulting in a total of 3840 stimulus presentations per session). Each block began with the presentation of a target orientation (drawn at random, without replacement, from the 16 stimulus orientations), displayed centrally as a green line (4° length). The stimulus stream consisted of randomly oriented Gabor patches, presented centrally for 100 ms, at an average rate of 650 ms. Stimuli had 16 possible angles (5.625-174.375°, in steps of 11.25°). Participants were instructed to respond whenever a Gabor patch with a matching orientation appeared. Since stimuli were drawn uniformly from the 16 possible orientations, 1/16 of all stimuli were targets. The angles were encoded into two covariates using the sine and cosine functions. Each block was cut into three shorter segments, giving participants brief rest periods. During the rest periods, the target orientation was presented again as a reminder. Participants were instructed to respond as quickly and accurately as possible.

### MEG data acquisition and preprocessing

Neuromagnetic data were acquired using a whole-head VectorView system (204 planar gradiometers, 102 magnetometers, Elekta Neuromag). The signals were sampled at a rate of 1000 Hz and on-line band-pass filtered between 0.03 and 300 Hz. Data were preprocessed using the OSL software library^1^. The raw MEG data were visually inspected for artefacts, de-noised and motion-corrected using Maxfilter Signal Space Separation (Taulu et al., 2004), and downsampled to 250 Hz. Artefacts arising from eye blinks and heartbeats were removed via independent component analysis. Epochs were generated around each stimulus onset (from 0s to 0.6s) and visually inspected to eliminate any remaining trials with excessive noise.

### Standard decoding analysis

The standard approach for decoding, illustrated in **Figure 1a**, estimates one decoding model at each time point. We will assume for simplicity that the stimulus is a continuous variable, such that a decoding model is a regression model (in this study, the stimulus is represented by two continuous variables or features: sine and cosine of the corresponding angle). For a categorical variable (e.g. the type of stimuli), the equations below can be easily adapted to use, for instance, logistic regression. Further extensions using more complex estimation methods (support vector machines, neural networks, etc.) are also possible if they are formulated within the Bayesian paradigm.

Let *t* = *1…T* index time, let *X_t_* be a (trials by channels) matrix containing the data at *t*, and let *y_t_* be a (trials by stimulus features) vector containing the stimulus. The solution for the decoding model at *t* is computed as

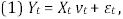

where *ε_t_* is Gaussian-distributed noise and *v_t_* is a (channels by stimuli) matrix of decoding weights. Given this, *v_t_* is typically obtained by maximum likelihood as

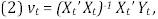

where’ represents matrix transposition. In this case, we decode the pair of variables formed by the sine and cosine of the angle of interest, such that *Y_t_* has two columns. In this paper, also, the data was projected into 48 principal components (explaining on average 96% of the variance in the data) for computational reasons, such that *v_t_* has dimension (48 by 2).

### Temporally Unconstrained Decoding Analysis

We introduce a novel probabilistic model containing each of the decoding models and the probability of each decoding model to be active at each time point at any given trial; also, the model includes a transition probability matrix, indicating the probability of transitioning from one decoding model to another within the trials. The proposed approach is illustrated in **Figure 1b** (right). In this case, instead of *T* different decoding models, we estimate only *K* decoding models (*m_k_*) where each corresponds to a (channels by stimulus features) matrix of decoding weights *w_k_*. (As before, the data was projected into 48 principal components and the number of stimulus features is two, such that *w_k_* has dimension 48 by 2.) Given time point *t* and trial *s*, let us define *y_st_* as the (1 by stimuli) value of the stimulus, and *X_st_* as the (1 by channels or PCA components) data vector. The top-level model in this case is formulated as

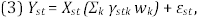

where *γ_stk_* = *Pr_st_(m_k_)* is the probability of model k being active at time point *t* and trial *s*, and *ε_st_* is Gaussian-distributed noise at time point *t* and trial *s* (with a shared variance that is not model-dependent). Note that, whereas the decoding weights are defined at the group level, the probability of model *k* being active at time point is specific of each trial, and, therefore, the decoding dynamics are allowed to be also trial-specific. Furthermore, we model the probability of transition between decoding models as

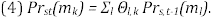

This way, the probability of a decoding model to be active at some time point depends not only on its decoding performance but also on the decoding model that was active in the previous instant for this trial. Besides the transition probabilities *Θ_l,k_*, we also model the initial model probabilities *π_k_*, referring to which is the model active at the start of the trials.

### Inference of the parameters

Given the data, we need to estimate the weights *w_k_*, the probabilities *γ_stk_*, the transition probabilities *Θ_l,k_*, and the initial probabilities *π_k_*. Here, we perform the estimation in two separate steps: first, the decoding weights, and then the other parameters. Although it is possible to perform the estimation of all parameters simultaneously, we adopt this strategy so that the results are comparable to the standard approach.

In order to estimate *w_k_*, we take a start with the standard decoding approach described above by estimating one decoding model *v_t_* at each time point, such that all trials all pooled together at each time point. We then compute the error of each model *v_t_* for each time point as

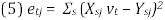

In words, *e_tj_* is the across-trials error of model *v_t_* evaluated at time point *j*. For each pair of time points *(i,j)*, a measure of divergence between the corresponding models *v_i_* and *v_j_* can be then obtained as

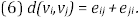

Based on *d(v_i_,v_j_)*, we use hierarchical clustering to group the *T* decoding models into *K* clusters; the representatives of these clusters constitute our first approximation to the decoding models *w_k_*. Note that these models are based on the standard approach and are constrained to be synchronous across trials, i.e. at a given time point the same decoding model is active for all trials. Still keeping this restriction, we then refined the estimation of *w_k_* by using the expectation-maximisation (EM) algorithm, where we alternatively estimate *w_k_* and the time points when *w_k_* is active (i.e. the EM algorithm is initialised with the cluster representatives from the previous step). At this stage, we stress, if *w_k_* is active at time point *t* that means that it is active for all trials at this time point. In the Results, we referred to having *K* instead of *T* decoding models, while still restricting the decoding models to be synchronous. This corresponds to the output of this step, which constitutes the initialisation for the next step.

In the next step, then, we fix the estimation of *w_k_* to the previous estimation and proceed to dispense with the synchrony restriction. For this, we use a Bayesian approach, estimating the a posteriori distribution of *γ* and *Θ* using variational inference (Wainwright and Jordan, 2008). The estimation of these parameters corresponds to the Hidden Markov model (HMM) forward-backward equations, described elsewhere (Rabiner, 1989; Vidaurre et al., 2016). For this step, we reused the equations from the HMM-MAR model (where MAR stands for multivariate autoregressive model; Vidaurre et al., 2016) and functions from the corresponding toolbox^2^. More specifically, we fed both the data and the stimulus to the model, and used a restricted MAR model of order 1 (Vidaurre et al., 2016), such that the autoregressive coefficients that predict the data as well as the autoregressive coefficients that predict the stimulus using the previous value of the stimulus were not modelled (i.e. the only autoregressive coefficients to be modelled are the ones that predict the stimulus from the data). We then delayed the data one time point with respect to the stimulus, such that the stimulus at time point *t* gets predicted by the data at time point *t* (instead of *t-1*); for specific implementation details, we refer to the online documentation of the toolbox^3^.

### Predicting reaction time

Unlike the information of the stimulus itself, the information of reaction time (RT) was not included in the model. That allowed us to use the relation of such model time courses to RT in order to add further confidence on the biological relevance of the estimated models. For this, we used model time courses to predict RT, discarding those trials where a button press was not effected. More specifically, we used principal component analysis to reduce the regressor dimensionality from *T* (no. of time points) per trial to 25 principal components. This was done for each decoding model separately. We then used sparse regularised regression (Vidaurre et al., 2013) where the regularisation parameter was itself chosen using cross-validation

### Encoding models

As discussed in (Weichwald et al., 2015), interpreting the magnitude of the decoding weights *w_k_* is not straightforward. If we wish to examine the spatial extents of the regions involved in stimulus processing, we need to use encoding models as in (Myers et al., 2015). That is, instead of using weights that predict the stimulus using the data from the entire sensor-space, we construct spatial maps using the encoding weights *b_lk_* that, from the stimulus, predict the data separately at each sensor *l*. By using the decoding model time courses *γ_stk_*, it is straightforward to associate a set of encoding weights *b_lk_* to each decoding weights matrix *w_k_*. The encoding weights are computed as

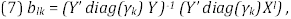

where *X^l^* represents the concatenated data for sensor *l*, *γ_k_* are the concatenated decoding model time courses for state *k*, and *diag()* diagonalises the vector argument into a diagonal matrix. In this case, *b_lk_* has two elements because *Y* has two columns (sine and cosine of the corresponding angle). We finally use the *b_lk_* weights to obtain the explained variance per sensor as shown in **Figure 4**.

### Cross-validation

One important question investigated through decoding is when (and where) the brain is processing the stimulus. This is typically addressed by using cross-validation, where, for each cross-validation fold, we estimate a decoding model at each time point using the training trials, and test each of the models in the held-out trials at each time point. For TUDA, as discussed above, this is problematic because the temporal information of the decoding models in the held-out trials is unknown. If we are to use cross-validation to compare different models (for instance, the standard decoding approach and TUDA), either we cannot use the stimulus information, or else we need to correct for the bias brought about by using the model time courses in the held-out trials. Here, we do the latter and correct the introduced bias by the use of surrogate data.

We generated random samples (surrogates) of the data set by permuting the labels (here, the angle values) across trials. We then run cross-validation on each surrogate data set, and computed the 5% percentile of accuracy across surrogates. These provided, for each cross-validation approach, a baseline that we can then subtract from the original cross-validation estimates. We refer to the result as *Adjusted CV-R^2^*. Since we are now comparing differences to the surrogate accuracies, overfitting gets accounted for and we can then compare between different approaches including the standard decoding. Note that we carried out this procedure for TUDA in two different ways: either using the model time courses estimated on the entire data (i.e. considering between-trial temporal differences in the held-out data), or by using the model that is most active on average at each time point in the held-out trials (losing this temporal variability).

## Discussion

In this paper, we use the Hidden Markov Modelling framework to propose a new approach capable of *time-resolved decoding*, referred to as Temporally Unconstrained Decoding Analysis or TUDA. Using TUDA, it is possible to bypass the highly constraining assumption of consistent timing of states (or decoding models) across trials as made by the traditional decoding approach. By making decoding temporally unconstrained, we are able to gain new insight into the nature, sequence, and temporal variability of neural states that contribute to the perceptual judgements made on different experimental trials.

An increasing number of studies have used time-specific pattern classification methods to show that stimulus-specific patterns are highly time-specific and do not cross-generalize across time points. Further, these methods suggest that neural processing traverses a large number of states in a smooth, stereotyped cascade (King and Dehaene, 2014). This could occur through transitions from one attractor state to another within one circuit (Miller, 2016; Durstewitz et al., 2010), or, similarly, through sudden transitions from a null state to a stimulus-specific state in a downstream brain area (Latimer et al., 2015). However, this trajectory through neural state space assumes that trajectories are highly reproducible across trials. By relaxing this assumption, we found that processing might possibly have slower dynamics than suggested by the standard decoding approach. Importantly, we found these dynamics to be related to behaviour, exemplified here as reaction time. We next discuss some additional aspects about methodology and interpretation.

### Alternative decoding models

Here, we have used the instantaneous (raw) sensor-space MEG signal to predict the stimulus, in analogy to the study from Myers et al (2015) based on the same data set. Both Myers and colleagues’ approach and ours use information about the relative magnitude between sensors in order to decode, and disregards other information. Given that the stimulus at time point *t* is predicted using only data at time point *t*, this approach is, for example, blind to the oscillations that encompass the instantaneous signal used for the prediction.

More powerful (or interpretable) extensions, where the phase of ongoing oscillations is used for the prediction, are straightforward to implement under the proposed HMM-based framework. For instance, similar to Vidaurre et al. (2017c), the signal can be “embedded” such that a window around *t* (and not only t) is used to predict the stimulus at time *t*, effectively incorporating phase information into the estimation. Other possibilities are to use information of phase only (Cabral et al., 2017) or power only (Baker et al., 2014). Unlike the embedded approach, which uses the raw signal without the need of any mathematical transformation, these alternatives are based on the Hilbert transform and the use of filtering, for which an ad-hoc selection of the frequency bands of interest is required. Besides the decision of which features of the data will constitute the base for the prediction, another possible extension is to replace the simple, linear regression model by more powerful prediction algorithms such as support vector machines or neural networks (Hastie et al., 2001) as far as these are formulated within the Bayesian framework. This scenario, where the decoding models may have a much larger number of parameters, can easily be handled with the proposed framework (where there are only *K* models) but is less manageable for the standard approach (where there are *T* models), especially if the number of trials is not very large.

### Decoding with fMRI

In this work, we found temporal variability between trials in the range of hundreds of milliseconds. Although this is a significant amount of time when considering electrophysiological data, fMRI has much lower temporal resolution, and the temporal uncertainty brought about by the hemodynamic response further hampers the benefits of our approach for probing tasks with fast cognitive mechanisms (attention, perception, etc.). There are however tasks with meaningful variability at the fMRI scale (seconds rather than milliseconds): difficult decision-making, mind-wandering, tasks with components that fluctuate slowly such as arousal, and tasks with delayed activity such as those related to working memory. In these types of tasks, the proposed method has the potential to excel at discovering temporal variability, up to the limit imposed by the modality’s inherent temporal resolution and its haemodynamics.

### Localised decoders

Here, we applied the model to whole-brain sensor space data, in line with previous work (Myers et al., 2015). Finer spatial and temporal information can be obtained from applying this method to source-localised data, possibly running the model on one group of regions at a time. This approach can give us insight on the different temporal dynamics of various regions in encoding the stimulus by examining and post-hoc comparing the model time courses between regions. This strategy would be comparable to the analyses performed by Baldassano et al. (2017) with an unsupervised HMM (i.e. trained with no information of the task), where sequences of states where estimated from different brain regions while subjects watched movie-based stimuli. This study revealed that higher-order regions follow state segmentations that match the movie structure more closely than those followed by sensory regions. By including the stimulus (or certain aspects of it) into the model, we can however target more specific aspects of cognition and will benefit from higher sensitivity.

### Null states

The HMM is a general framework that has been used previously to describe brain activity in an unsupervised fashion (see e.g. Engel et al., 2016; Baker et al., 2014; Vidaurre et al., 2016; Vidaurre et al., 2017a; Vidaurre et al., 2017b; Vidaurre et al., 2017c; Baldassano et al., 2017). Here, we draw from this general framework to handle the supervised setting, where each HMM state corresponds to a certain particular relationship between bran activity and the stimuli. But, what happens for these time points where there is no relationship between brain activity and the stimulus at all (for instance, because the brain has not yet encoded the information in any way)? In an unsupervised setting, because the brain is never silent, all states are always meaningful (they always represent something, because there is brain activity in all time points). In a supervised setting, however, it is useful to detect when there is *nothing* to represent. In the current implementation, the proposed model can express this circumstance by using a “null” state (where the decoding weights are close to zero), or, instead, by using some random mixture of (otherwise meaningful) states such that, at these “empty” time points, the decoding error is higher than when these states are faithfully representing the stimulus. Post-hoc analyses would be required to detect this situation. A potentially more useful strategy would be to fix one decoding state to be the null state (i.e. having fixed, zeroed decoding weights), such that it will become active when the examined regions are unaware of the stimulus. In the data used in this paper, nevertheless, this would likely be of little use given the short inter-stimuli intervals.

## Conclusion

In this paper, we proposed a novel method for neural decoding, referred to as TUDA, where we dispense altogether with the assumption that neural processing is timed consistently across trials. Our results, on a simple perceptual decision-making task, indeed suggest that the assumption of consistent timing over trials made by a traditional decoding approach is not always justified, and can lead to misinterpretations of the dynamics of the cognitive mechanisms underpinning the processing of the stimulus. Although we have focused on a relatively simple stimulus, the technique can straightforwardly be applied to more complex cognitive tasks including volitional behaviour. Our approach also makes it possible to analyse the between-trial temporal variability, which, as shown above, can hold a significant relationship to behaviour, and could correspond to changes in attention or to plasticity.

## Acknowledgements

This study was supported by the NIHR Oxford Health Biomedical Research Centre, an MRC UK MEG Partnership Grant (MR/K005464/1), a James S. McDonnell Foundation Understanding Human Cognition Collaborative Award (220020448), and the NIHR Oxford Health Biomedical Research Centre. The Wellcome Centre for Integrative Neuroimaging is supported by core funding from the Wellcome Trust (203139/Z/16/Z). DV is supported by a Wellcome Trust Strategic Award (098369/Z/12/ Z). MWW is supported by the Wellcome Trust (106183/Z/14/Z) and the MRC UK MEG Partnership Grant (MR/K005464/1). ACN is supported by a Wellcome Trust Senior Investigator Award (ACN) 104571/Z/14/Z.

## Supplemental Information

We generated synthetic data in order to demonstrate the basic functioning of TUDA, and to show, in a scenario were the ground-truth is known, how the standard decoding approach can incorrectly suggest a large number of information processing states and a rapid succession of state changes. We describe this setting here in detail.

### Synthetic experimental design

We sampled N=1000 trials of one second duration, assuming sampling frequency of 300Hz (i.e. 300 time points per trial). The stimulus is modelled to be a colour, with three features representing RGB coordinates. For each trial, the colour stimulus was randomly sampled from a 3-dimensional uniform distribution (with values between 0 and 1). The data is set to have twelve channels. We assume there are three “cognitive processes” (or ground-truth states) underlying stimulus processing. Each of these is modelled as a (3 RGB coordinates by 12 channels) matrix of coefficients. At each time point and trial, the data are generated by multiplying the current colour (1 time point by 3 RGB coordinates) by the corresponding matrix of coefficients (3 RGB coordinates by 12 channels), and then adding some Gaussian noise (standard deviation equal to 0.1). We set each of the ground-truth states to activate or deactivate a different subset of the channels (4 channels per state). More specifically, for each state we sample the (3 RGB coordinates by 4 channels) active coefficients from a uniform distribution, and the rest are set to zero. All trials are set to start in state 1, and transitions are always from state 1 to state 2, from state 2 to state 3, and from state 3 to state 1. The duration of the state visits is sampled from a uniform distribution ranged in between 0.066s (20 time points) and 0.166s (50 time points), such that each state is typically visited three times per trial.

### TUDA discovers the ground-truth states

Assuming the right number of states (three), we ran the TUDA inference in order to estimate the decoding model time courses as well as the decoding coefficients and the transition probabilities between models. **Figure 2a** shows, on top, the average model time courses (which can be interpreted as the probability to be in each state or model at each time point) and the model time courses for one example trial; at the bottom, it is presented the estimated model time courses, on average and for one trial. It can be observed that the model inference is accurate both in terms of the average and for the single trial.

### Standard decoding overestimates the number of processing states

Using the standard approach for decoding, it has often been observed that the decoding models (e.g. the regression weights) continuously fluctuate as a function of time. This sometimes leads to poor decoding generalisation across time - i.e. a model trained at one time point has low accuracy when tested at a different time point (King and Dehaene, 2014). If we consider such regression weights as a proxy of the underlying neural processes that are relevant to the task, it can be thus inferred that such brain processes are highly dynamic (see e.g. Meyers et al., 2008; Carlson et al., 2011; Isik et al., 2013; Carlson et al., 2013; Stokes et al., 2013; King et al., 2014); that is, the brain goes through a large number of different processing states that are not necessarily interchangeable. Here we argue that this can potentially be caused in an artefactual manner by the temporal variability between trials. We ran standard decoding on our simulated data (where there are only three states with an average dwelling time of 0.11s) to obtain, by pooling across trials, one model per time point. Each of these models have 12×3=36 regression coefficients. We then computed the Pearson correlation between sets of regression coefficients for each pair of models and ran hierarchical clustering. The results are shown in **Figure 2b**, where we can see a large fragmentation of models (the models are not ordered by time here, but were automatically ordered seeking spatial contiguity). Following the approach of assessing the generalisation across time in a cross-validation setting (King and Dehaene, 2014), we tested the model estimated at each time point on the held-out trials at the entire range of time points. **Figure 2c** shows the corresponding generalisation matrix, using CV-R^2^ as the summarising statistic. It can be observed that the trials start sharply as state 1 is always active at the beginning of the trial. Then, the pattern becomes blurrier as a consequence of the increasing between-trials temporal variability.

---

**Figure SI-1.**
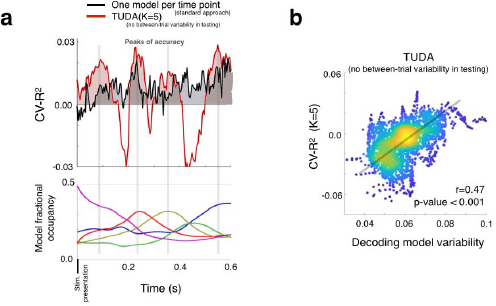
Cross-validation underestimates TUDA’s performance if we lose between-trials temporal variability in the held-out trials. **(a)** For the same illustrative session used in **Figure 2**, CV-R^2^ is shown as a function of time for the standard approach (black) and the proposed model when using *K*=5 decoding models (red). Underneath, the model fractional occupancy (see **Figure 2**) reveals that the peaks of accuracy closely correspond to the time points with less between-trials temporal variability. **(b)** There is a strong correlation between decoding model uncertainty (as expressed by the variability of the model time courses across trials) and accuracy (as expressed by the cross-validated explained variance, CV-R^2^), where each data-point in the scatter plot corresponds to a time-point within the trial (colour represents density of points and the line represents the slope of regressing model variability on CV-R^2^).

**Figure SI-2.**
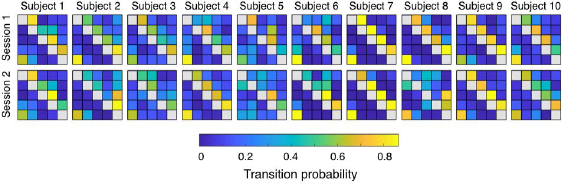
Transition probability matrices between decoding models for all sessions and subjects reveal that information processing follows consistent sequences of states in most of the sessions.

## Footnotes

1 https://ohba-analysis.github.io/osl-docs

2 https://github.com/OHBA-analysis/HMM-MAR

3 https://github.com/OHBA-analysis/HMM-MAR/wiki/User-Guide#restrict

